# starbase: A Database and Toolkit for Exploring Giant Cargo-Carrying Mobile Elements in Fungi

**DOI:** 10.64898/2025.12.04.686951

**Authors:** Adrian Forsythe, Emile Gluck-Thaler, Aaron Vogan

## Abstract

*Starships* are a recently discovered superfamily of extremely large mobile genetic element (MGE)s in fungi that encode diverse gene sequences, many of unknown function. *Starships* are widespread throughout filamentous Ascomycetes (Pezizomycotina), but relatively little is known about their fine-grained distributions at lower taxonomic levels. As more and more *Starships* are discovered, it is increasingly important to more effectively catalog their gene contents and taxonomic distributions to better understand their contributions to fungal evolution. To address this, we developed `starbase`, a web server and comparative toolkit for exploring *Starship* diversity and hypothesis generation. The `starbase` database is constructed from *Starships* identified from existing studies, as well as an exhaustive *de novo* survey of *Starship* sequences within a set of 19 863 publicly available fungal genome assemblies. This database consists of 5 493 *Starships*, their associated nucleotide sequences, captain gene protein sequences, cargo gene annotations, and other metadata pertaining to the annotation and analysis of *Starships* in fungal genomes. As a resource, `starbase` provides new avenues for studying structural variation in fungal genomes. `starbase` provides several key features for the research community: a centralized repository of curated *Starship* annotations, a standardized accessioning system enabling consistent referencing of elements across studies, tools for searching existing sequences and classifying novel *Starships* based on established classification schemes, and a submission portal encouraging community contributions. As *Starship* identification becomes a routine component of fungal genome annotation, `starbase` provides a framework for organizing this growing body of data and facilitating comparative analyses across the expanding landscape of fungal genomic diversity.

## 1 Background

### 1.1 Introduction to MGEs

Mobile genetic elements (MGEs) are found throughout the tree of life [10] and are typically present in variable quantity and diversity within individual species [11–13]. In microbial genomes, MGEs mobilize substantial portions of their microbial hosts’ genetic material, including genes involved in pathogenicity, metabolism, and coping with environmental stressors [14–18], thus playing a crucial role in genome evolution and adaptation [19–21]. MGEs that mobilize one or more genes with functions beyond transposition and transfer are known as cargo mobilizing elements (CMEs). In addition to nucleic acid mobilization, the genes carried by CMEs encode diverse molecular functions. Predominantly studied in bacteria, eukaryotic CMEs have increasingly been the subject of recent research [22–26], including the description of a new superfamily of transposable elements, the *Starships* [1–3]. *Starships* are endemic to the fungal subphylum Pezizomycotina and are extremely large, ranging from approximately 20 to 700 kb in size. *Starships* are mobilized by a tyrosine recombinase (YR) gene, referred to as the “*captain*” [3]. In contrast to many smaller MGEs, *Starships* can be identified based on sequence motifs, which define the element boundaries. These include short direct repeats (DRs) at the termini of the elements and often short asymmetric tandem inverted repeats (TIRs) [27]. In addition to carrying genes for their own mobilization, they also carry diverse sets of cargo genes, which generates an extreme amount of variation among elements [7].

Initially, *Starship* sequences were manually annotated based on comparative genomic analyses performed on select collections of fungal genomes. Following these studies, automated methods for discovery and curation of *Starship* sequences were developed and applied to expansive collections of fungal genomes, uncovering a large number of diverse elements [1, 7]. To organize the diversity of *Starships* a classification scheme has been proposed whereby elements are grouped at three hierarchical levels. The first is family, based on monophyletic clades within a phylogeny of the YR domain of the *captains*. The second is navis (plural naves), again based on comparisons of *captain* sequences, but defined by amino acid-based orthogroups. The third and final is haplotype, based on k-mer similarities across the entire element nucleotide sequence.

While this provides a general framework for organizing and comparing *Starship* sequences, precise values for cutoffs at each classification rank are not employed, providing some flexibility in naming elements. However, this approach also complicates approaches to catalogue elements at scale.

Although several databases currently exist to catalogue MGEs [28–33], they are mostly focused either on bacterial MGEs, or smaller eukaryotic transposable elements (e.g. repbase, Dfam). Furthermore, existing databases of MGEs are often built without a comprehensive classification schemes or ontology information, making it difficult to track and analyze these elements systematically across species. *Starships* are distinguished from other MGEs, by their size, capacity to carry a wide repertoire of cargo genes [34], and usual presence in a single copy within the genome (with some exceptions). As a result of these factors, the construction of consensus sequences (common within other MGE databases) is not meaningful or indeed feasible. Similarly, the creation of HMMs for classification, a requirement for the Dfam database [35], is also difficult and infeasible given the amount of sequence diversity present between *captain* genes. These factors highlight the challenges for integrating *Starships* into existing databases of MGEs/CMEs.

Phylogenetic distributions, bioinformatic analyses, and laboratory experiments all suggest that the *Starships* move not only within genomes, but between them, serving as a mechanism for horizontal gene transfer (HGT) in fungi [1, 3, 34, 36, 37]. It is thus becoming clear that *Starships* play a substantial role in facilitating rapid adaptation through the transfer of multiple genes in a single transfer event [1, 38]. Additionally, the number of identified *Starship* sequences has grown rapidly (Table 1) since the first description of *Starships* in [2]. For these reasons, we believe that the development of a framework capable of maintaining and growing the information related to *Starships*, their host genomes, and their genetic cargo is crucial for future investigations of fungal genome evolution and adaptation. To address this, we have developed `starbase`, a comprehensive database and toolkit for the identification, classification, and exploration of *Starship* elements. The goal of `starbase` is to provide a publicly accessible resource of existing *Starship* sequences with curated annotations with standardized classification, all while adhering to FAIR (Findability Accessibility Interoperability and Reusability) principles [39]. This database consists of existing *Starship* sequences from previous studies, as well as the results of a large-scale genomic survey conducted as part of the database release, to identify similar *Starship* sequences in fungal genomes. In addition to the database, `starbase` serves as a toolkit for researchers, enabling the identification, classification, and comparative analysis of *Starship* elements. `starbase` was developed as a resource for the community, and provides an opportunity for researchers interested in *Starship* biology to submit candidate *Starship* sequences to be included in successive database releases, pending review. This breadth of diversity captured by the sequences in `starbase` demonstrates the wide-reaching impact and potential of this resource for fungal genomics.

**Table 1.** Summary of the main sources of annotated *Starship* sequences used to build starbase.

| Type of Search | Citation | Number of Genomes | Number of *Starships* |
| --- | --- | --- | --- |
| Manual curation | [1–4] and this study | ~100 | 169 |
|  | [5] | 59 | 54 |
|  | [6] | 1 | 1 |
| *Starship* annotation with **starfish** | [7] | 1 148 | 1 148 |
|  | [8] | 1 455 | 1 626 |
|  | [9] | 29 | 29 |
| *Starship* identification with **minimap2** | This study | 19 863 | 2 466 |
| Total Number of *Starships* in **starbase** |  |  | 5 493 |

## 2 Construction and Content

### 2.1 Framework and Core Features

`starbase` is a web application built in Python 3.9 using `plotly Dash` [40, 41], which extends `python Flask` [42] to provide reactive web components. The application employs a client-server architecture with a SQLite relational database backend for data management, with database interactions managed through `SQLAlchemy` [43] engine and sessions. The front-end is built using Dash Bootstrap [44] and Dash-Mantine [45] components, providing interactive and responsive user interface elements. Data visualization capabilities are implemented through `plotly`, `blasterjs` [46] for BLAST result visualization, and `clinker`/`clustermap.js` [47] for comparative genomic visualization. The complete application is packaged as a Docker image and deployed to the SciLifeLab Serve platform, a hosting service maintained by SciLifeLab (Uppsala, Sweden), which serves the application within a kubernetes pod. `starbase` is accessible at starbase.serve.scilifelab.se.

Through the web interface, users can browse information for existing *Starships*, download *Starship* sequences, search and classify their own sequences against databases of existing *Starship* and *captain* sequences, visualize and compare genomic features of *Starships*, and submit sequences for future curation. These capabilities are organized across several functional modules. The wiki page provides searching and browsing functionality for existing *Starships*, where users can filter elements using multiple criteria: i) taxonomy of the host genome, ii) *Starship* family classifications, and iii) specific `starbase` Ship Accessions (SSAs). The taxonomic distribution of genomes harboring *Starships* is visualized via a hierarchical sunburst chart that updates dynamically with applied filters. For each *Starship* family, we provide summary characteristics including the number of elements, minimum and maximum length, publication information for previously annotated *Starships*, and sequence logo plots of direct repeats identified in curated *Starships*.

The sequence search module enables users to classify their own sequences against databases of existing *Starship* and *captain* sequences. A `BLAST` search can be initiated using either a nucleotide or protein query sequence. If a nucleotide sequence is provided, a `blastn` search is conducted against a database of full *Starship* sequences, and the protein sequence of the *captain* gene is extracted using DIAMOND [48]. The extracted *captain* protein sequence (or a user-provided protein sequence, if that was the initial query) is then compared against a HMM profile of *captain* genes `hmmsearch` [49] to identify the closest *Starship* family present in `starbase`. `BLAST` results are visualized using the `blasterjs` library and can be exported in TSV format. Users can also visualize and compare genomic features of *Starships* using the synteny viewer, which is powered by `clinker`/`clustermap.js`.

### 2.2 Database Curation and Annotation Pipeline

To organize and maintain *Starship* metadata and gene annotations, we adapted an existing framework developed for plasmid sequence curation and management, Manual Annotation Studio (`MAS`) [46, 50], and modified it for *Starship* sequences. The extended `MAS` framework includes an automated gene prediction pipeline and tools for manual curation of sequences and meta-data, enabling standardization of gene annotations across *Starships*. The annotation pipeline implemented in `MAS` is particularly appropriate for a database of rapidly expanding sequences, as gene annotations and their associated metadata are updated based on similarity to existing sequences already in the database. As new sequences are submitted, automated gene discovery and annotation is performed using several reference databases, with the output visualized for manual curation and review through the web interface.

We maintain two databases: the public `starbase` database and an internal `MAS` database, which represent the public and private sides of *Starship* curation and annotation, respectively (Figure 1A). The `starbase` database is organized around a relational schema encompassing *Starship* sequences, associated metadata, genomic features, publication references, and gene annotations (Table 1C). This includes taxonomic classifications of host genomes, *Starship* classification information (family, navis, haplotype), detailed information about the fungal host genome assembly, structural features (TIRs, DRs, and insertion sites), and cargo gene annotations.

**Figure 1.**
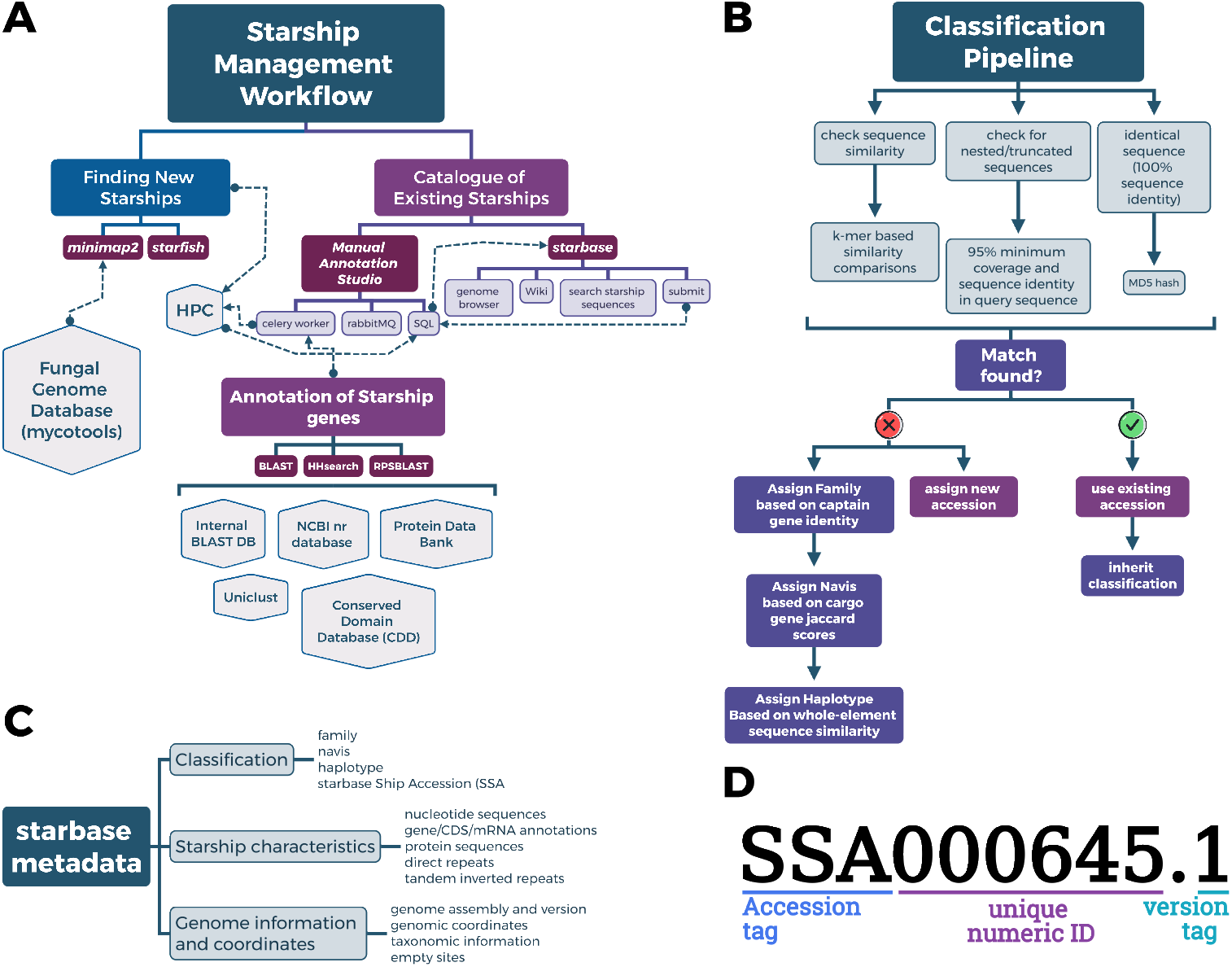
A) Overview of the `starbase` architecture showing the relationship between the public-facing database, `starbase`, and the internal system for *Starship* annotation (`MAS`). Database components are shown with their interactions and information flow. B) Classification workflow implemented in `starbase`, illustrating the stepwise process for identifying and classifying new *Starship* sequences based on existing database entries and sequence characteristics. C) `starbase` Ship Accessions (SSA) provide a standardized nomenclature for *Starship* identification. Any given SSA consists of a unique six-digit numerical identifier and version suffix. These accessions provide a system for identifying any unique *Starship* sequence, with any completely identical (or nested) *Starship* sequences assigned the same SSA.

The internal `MAS` database architecture is based on the existing `MAS` project [50], which maintains a relational schema in `mariaDB` encompassing all information contained in `starbase`, as well as uncurated *Starships* and their gene annotations. `MAS` uses the `luigi` pipeline frame-work and celery task queue for managing computational tasks related to gene annotation within *Starships*. Genes are annotated using `BLASTP` [51] (against SwissProt [52], NCBI NR [53], and an internal database of *Starship* sequences), HHSearch [54] (against PDB [55]), `RPSBlast` [51] (against Conserved Domain Database (CDD) [56]), and InterProScan [57]. Automatic validation of protein sequences includes checking for valid start codons and amino acid characters. `MAS` also implements consensus-based annotation updates by automatically standardizing annotations based on similar sequences, with automatic rebuilding of internal `BLAST` databases when new annotations or *Starship* sequences are added. The `MAS` database tracks historical changes to *Starship* metadata, classification, gene annotations, and curation activities by moderators. Manual validation of gene annotations incorporates the assignment of confidence flags by moderators to indicate the level of certainty for specific annotations.

We classify *Starships* into three curation status levels to maintain transparency about data quality. “Curated” *Starships* have complete annotations including identified captain genes and clear element boundaries with TIRs and DRs. “Intermediate” *Starships* are missing one of these criteria, while “uncurated” *Starships* lack all of these features but remain in the internal `MAS` database for ongoing analysis. This tiered approach allows us to present users with the most reliable data while continuing to develop annotations for less well-characterized elements. The public SQLite `starbase` database is generated periodically from the internal `MAS` database and includes only curated and intermediate status *Starships*. Released database versions are archived on Zenodo (`10.5281/zenodo.17533381`), ensuring reproducibility and providing access to historical database states.

### 2.3 Classification Workflow

In accordance with existing standards for *Starship* classification [7, 27], we implement a multistage sequence classification system for classifying new sequences (Figure 1B). This classification workflow incorporates a series of sequential steps: i) testing for exact matches using MD5 hash comparison, ii) checking for contained matches based on 95% minimum coverage and sequence identity from `minimap2` [58] alignment, and iii) searching for highly similar matches based on 90% k-mer similarity using `sourmash` [59] against sequences already in `starbase`. When a match to an existing *Starship* is found, the classification information is inherited from the existing entry. If no existing match is found, the classification workflow proceeds with assignment of *Starship* family (based on `hmmsearch` results or direct sequence comparison), *Starship* navis (based on *captain* sequence clustering), and haplotype (based on whole-sequence similarity using `sourmash` k-mer thresholds).

To maintain data integrity, we developed an accessioning framework within `starbase`. SSAs are assigned using a six-digit numerical code, providing a standardized, unique identifier system for *Starship* sequences (Figure 1D). Each SSA is used to represent a single or multiple *Starship* sequences which may be present in one or many genomes. *Starship* sequences are assigned a SSA based on sequence similarity, which can range from completely identical sequences (found in different genomes), sequences with slightly different boundaries (“nested” sequences), or those that are highly similar to existing sequences within `starbase` (Figure 1B). We also implement a versioning system similar to NCBI assembly accessions: when updates are made to a sequence, a numerical tag is appended to the accession. Accessions are permanently retained in the database and serve as a community resource, providing standardized nomenclature for sequence identification, improving reproducibility of research, and facilitating data sharing and integration with other databases.

### 2.4 Comprehensive Census of *Starships* Within Fungal Genomes

To determine the current extent of *Starship* diversity within existing fungal genomes, we surveyed a collection of fungal genome assemblies from NCBI. The management of genome sequences and annotations was facilitated using `mycotools` [60], a database framework specifically designed for organizing large collections of fungal genome sequences and annotations. In total, 19 863 fungal reference genomes were included in the analysis (an additional 18 715 genomes compared to [7]).

We developed a bioinformatics pipeline to identify matches to existing *Starship* sequences in genome assemblies (stand-alone pipeline available on Zenodo repository). This pipeline detects both complete and partial matches by mapping full *Starship* sequences to genomic contigs or scaffolds using `minimap2`. The `minimap2` aligner offers different presets for varying sequence divergence levels: `asm5` for approximately 0.1% divergence, `asm10` for approximately 1% divergence, and `asm20` for approximately 5% divergence. We applied customized alignment parameters including enhanced end-to-end alignment detection (end-bonus: 100), optimized gap penalties for both short and long gaps, and variable mismatch penalties to accommodate sequence variation. The pipeline identifies alignments occurring in the terminal regions (first and last 5000bp) of query sequences. When flanks are identified on the same contig or scaffold and are separated by a distance within the allowed maximum gap, the pipeline attempts to join them into a single sequence. This algorithm enables identification of both full-length elements (over 90% query coverage) and truncated elements (based on partial matches and pairing of flanking regions).

Through this systematic survey of fungal genomes, we incorporated an additional 2 466 *Starships* into `starbase`, representing a substantial expansion of the known diversity of these elements (Figure 2A). This comprehensive census enriches our understanding of *Starship* distribution across fungal lineages as these newly identified elements span diverse taxonomic groups within Pezizomycotina. By integrating these newly discovered sequences into `starbase`, we provide researchers with access to the most comprehensive collection of *Starship* elements assembled to date, enabling more robust comparative analyses and classification efforts.

**Figure 2.**
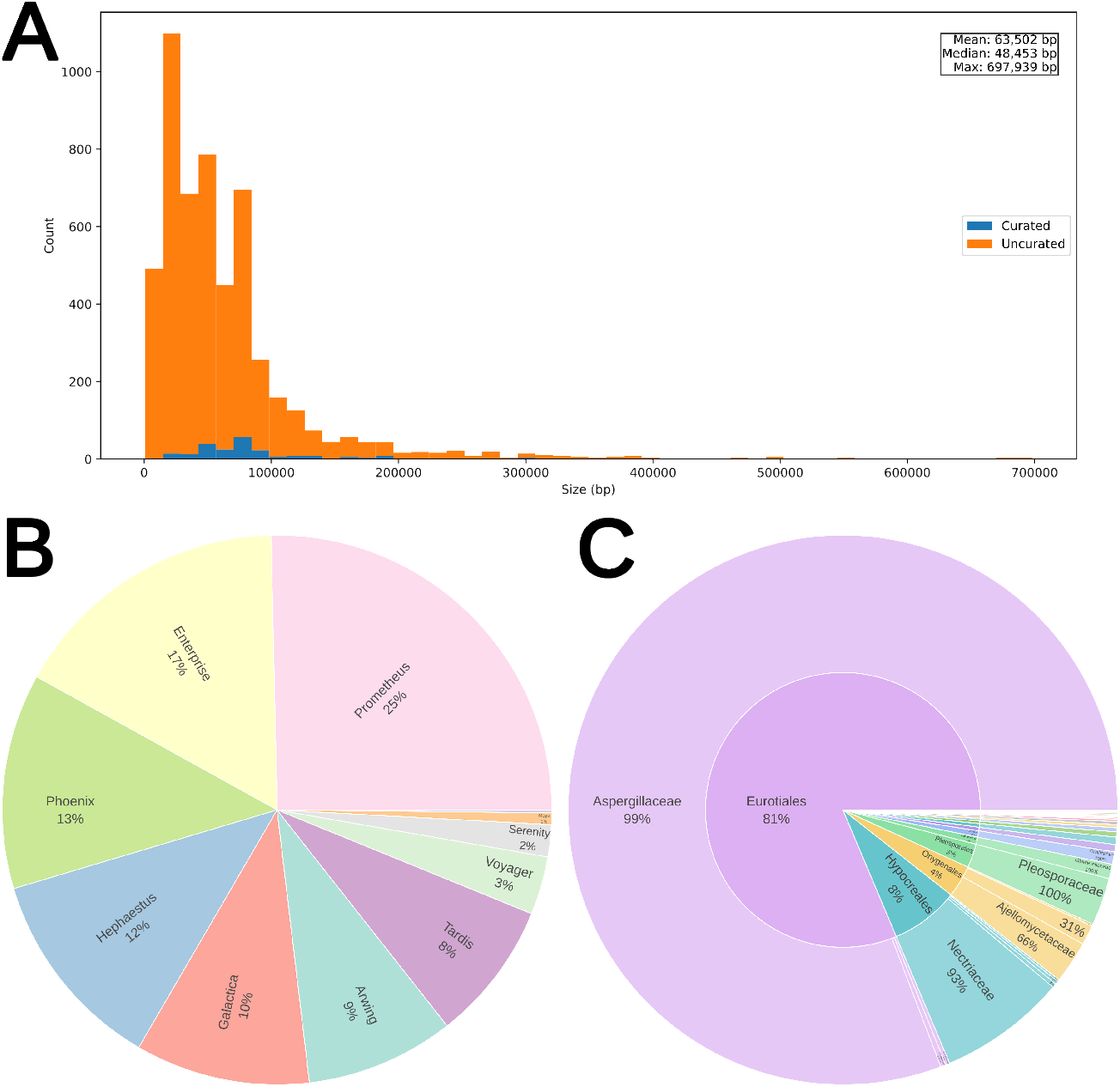
Summary of *Starships* in `starbase`. A) Distribution of *Starship* sequence lengths. B) Proportions of *Starships* classified within each *Starship* family. C) Taxonomic distribution at the Order and Family level of genomes harboring *Starships*.

### 2.5 Data Validation and Quality Control

Data integrity of `starbase` is maintained through a comprehensive validation pipeline designed to ensure accuracy and consistency. This multi-layered approach includes automated validation of *Starship* boundaries (TIRs, DRs), taxonomic information, and associated genome metadata. Motif recognition is used to identify potential boundary errors or assembly artifacts that might compromise element classification. Taxonomic assignments of fungal hosts undergo rigorous reconciliation against the NCBI taxonomy database [61], ensuring that host species information adheres to current nomenclature standards. Quality control extends to the gene annotation process through the confidence flagging system implemented in `MAS`. Version history tracking for all *Starship* sequences and their gene annotations enables tracing the provenance of information from initial identification through subsequent refinements. Database updates undergo systematic validation before release, with manual review of a representative sample of new submissions to verify correct classification and annotation. Version tracking ensures that all changes are documented and traceable, supporting reproducible research practices.

## 3 Utility and Discussion

### 3.1 Classification and Accessioning Framework

`starbase` provides both an accessible resource for exploring *Starship* diversity and a framework for ensuring consistency in the characterization of *Starships* across studies. The classification workflow implemented in `starbase` (Section 2.3) employs established criteria for *Starship* classification, enabling users to classify novel sequences based on these standards (for details see [7] and Box 2 in [27]). Together with this classification system, SSAs provide a stable reference point for *Starship* sequences, enabling reproducible investigations of specific elements across different fungal host genomes and genome assembly versions. Once assigned, SSAs remain fixed, with any updates (i.e. changes to element boundaries) tracked through incrementing versions (Figure 1D). As more *Starships* are discovered and submitted to the database, new SSAs can be assigned, ensuring that the nomenclature system grows alongside our knowledge of these elements.

### 3.2 Applications for Research

The comprehensive collection of *Starship* sequences in `starbase` enables several research applications in fungal genomics. First, the database facilitates comparative genomic analyses by providing standardized annotations and classifications across diverse *Starship* elements. This standardization allows researchers to identify conserved structural features and cargo genes, as well as lineage-specific adaptations that may reflect different evolutionary trajectories or ecological pressures. Second, the cataloging of *Starship* elements across fungal lineages enables investigation of HGT patterns. Researchers may track the distribution of closely related or identical elements across taxonomically distant hosts, providing evidence for potential horizontal transfer events. The combination of element-level similarity data (through k-mer analysis and sequence alignment) with phylogenetic context allows for evaluation of transfer frequency and the identification of fungal lineages that may be particularly prone to acquiring or donating *Starships*. Third, cargo gene annotations within *Starships* provide insights into the functional impact of these elements on host biology. By examining cargo gene repertoires across different *Starship* families and host lineages, researchers can investigate adaptive roles and evolutionary pressures shaping element sequence composition. The integration of structural features (TIRs, DRs, insertion sites) with sequence and annotation data enables comprehensive analyses ranging from mechanisms of transposition to patterns of cargo gene acquisition and loss. These analyses can reveal how *Starships* contribute to phenotypic diversity and adaptation in fungal populations.

### 3.3 Community Resource and Future Directions

`starbase` addresses a critical need in the fungal genomics community by providing a centralized, standardized repository for *Starship* sequences and metadata. The implementation of SSAs establishes a common nomenclature system, ensuring consistent referencing of elements across different studies and preventing confusion from ad-hoc naming schemes. This standardization becomes increasingly important as the number of characterized *Starships* continues to grow and their presence across diverse fungal lineages becomes more widely recognized.

The database architecture is designed with extensibility in mind, allowing for integration with other genomic resources and adaptation to accommodate new data types as our understanding of *Starships* evolves. Community contributions are facilitated through the submission portal, which provides a straightforward mechanism for researchers to contribute novel *Starship* sequences while maintaining data quality through the curation pipeline. As the database expands through community contributions and systematic genome surveys, `starbase` will serve as the primary reference for *Starship* research, supporting increasingly sophisticated comparative and evolutionary analyses of these remarkable mobile genetic elements.

## 4 Conclusion

The discovery and characterization of *Starships* represents a significant advancement in our understanding of fungal genome dynamics and evolution. These extremely large mobile genetic elements appear to be widespread within Pezizomycotina, with our comprehensive survey identifying an additional 2 466 *Starships* across diverse fungal lineages. The substantial variation in size, structure, and cargo gene content among these elements suggests diverse roles in adaptation and niche specialization among fungal species. The `starbase` database and toolkit provides resources for identifying, classifying, and analyzing *Starship* elements. By implementing standardized classification schemes and curated annotations, `starbase` establishes a foundation for consistent terminology and comparative analysis across different studies. The user-friendly interface and integrated visualization tools make these complex genomic elements accessible to researchers with varying levels of computational expertise, effectively lowering the barrier to entry for comparative analyses. The potential impact of *Starships* on fungal biology extends far beyond their role as interesting genomic features. Their capacity to mobilize extensive gene repertoires, including clusters relevant to metabolism, stress response, and virulence, suggests they may function as vectors for adaptive evolution. The identification of (near-)identical *Starship* sequences across fungal lineages can be used as the first indication of potential HGT events.

Looking forward, we anticipate that `starbase` will continue to evolve through regular database updates and expanded functionality. As research on fungal MGEs continues to advance, `starbase` will serve as a valuable community resource, promoting collaborative investigation of these fascinating genomic entities. Through this work, we hope to deepen our understanding of fungal genome evolution, the extent of HGT within fungi, and the contribution of *Starships* to adaptation/niche specialization for ecologically and economically relevant fungal species.

## Declarations

The authors declare no competing interests. A.A.V. was provided with support by funding from the Swedish Research Council VR (grant number 2021-04290) and ERC-2023-COG (Starship, 101126121) to A.A.V. Funded by the European Union. The source code for `starbase` is available at github.com/FungAGE/starbase. The release version of the `starbase` database is archived on Zenodo (`10.5281/zenodo.17533381`).

## Author Contributions

A.F. and A.A.V. developed the concept and general framework for `starbase`. A.F. is the developer of `starbase` and all related utilities and code, including the main collation of *Starship* data. All authors contributed to the annotation of *Starships* included in `starbase`.

## Acknowledgements

The authors acknowledge the resources provided by SciLifeLab (Uppsala, Sweden), including the SciLifeLab Serve platform, responsible for hosting of `starbase`. In addition, we thank Arnold Kochari, Jana Awada, Andrew Urquhart, and the members of the FungAGE Lab and Gluck-Thaler Labs for their help and feedback during `starbase` development.

## Acronyms

`BLAST`: Basic Local Alignment Search Tool
`MAS`: Manual Annotation Studio
CDD: Conserved Domain Database
CME: cargo mobilizing element
DR: direct repeat
FAIR: Findability Accessibility Interoperability and Reusability
HGT: horizontal gene transfer
HMM: Hidden Markov Model
MD5: Message-Digest Algorithm 5
MGE: mobile genetic element
NCBI: National Center for Biotechnology Information
SSA: `starbase` Ship Accession
TIR: tandem inverted repeat

